# [^18^F]FLT-PET as predictive non-invasive biomarker for neoadjuvant therapy with Wee1 and ATR inhibitors

**DOI:** 10.64898/2026.03.10.710900

**Authors:** Amirali B. Bukhari, Melinda Wuest, Frank Wuest, Armin M. Gamper

## Abstract

Besides immunotherapy, inhibitors of the DNA damage response (DDR) are currently one of the most promising contributors to improved cancer therapy. They exploit elevated replicative stress in cancer cells and often rely on synthetic lethality with existing gene deficiencies or between targeted pathways. In view of the absence of reliable histological biomarkers for replicative stress, this study examined [^18^F]-fluorothymidine (FLT) positron emission tomography (PET) as alternative or complementary approach to predict treatment response to DDR inhibitors. Using orthotopic and syngeneic triple negative breast cancer mouse models and treatment with combined AZD6738 and AZD1775 (inhibiting ATR and Wee1, respectively) this study found that: a) Sequential [^18^F]FLT-PET in the early phase of treatment was able to predict ATR/Wee1 inhibitor treatment efficacy, whereas b) [^18^F]FLT tumor uptake at onset of therapy was unable to predict treatment outcome, despite c) [^18^F]FLT tumor uptake positively correlating with Ki-67 staining, the clinically used proliferation marker. Importantly, non-invasive monitoring of changes in tumor biology by [^18^F]FLT-PET predicted which tumor model responds to combined AZD6738/AZD1775 treatment and established a quantitative correlation in [18F]FLT tumor uptake with tumor shrinkage in individual responders.

Since the inhibitors AZD6738 and AZD1775 are already in phase I/II clinical trials, this knowledge could soon be translated into the clinic. To our knowledge this is the first study to correlate non-invasive PET imaging with treatment efficacy of DDR inhibitors.

**Graphical Abstract:** 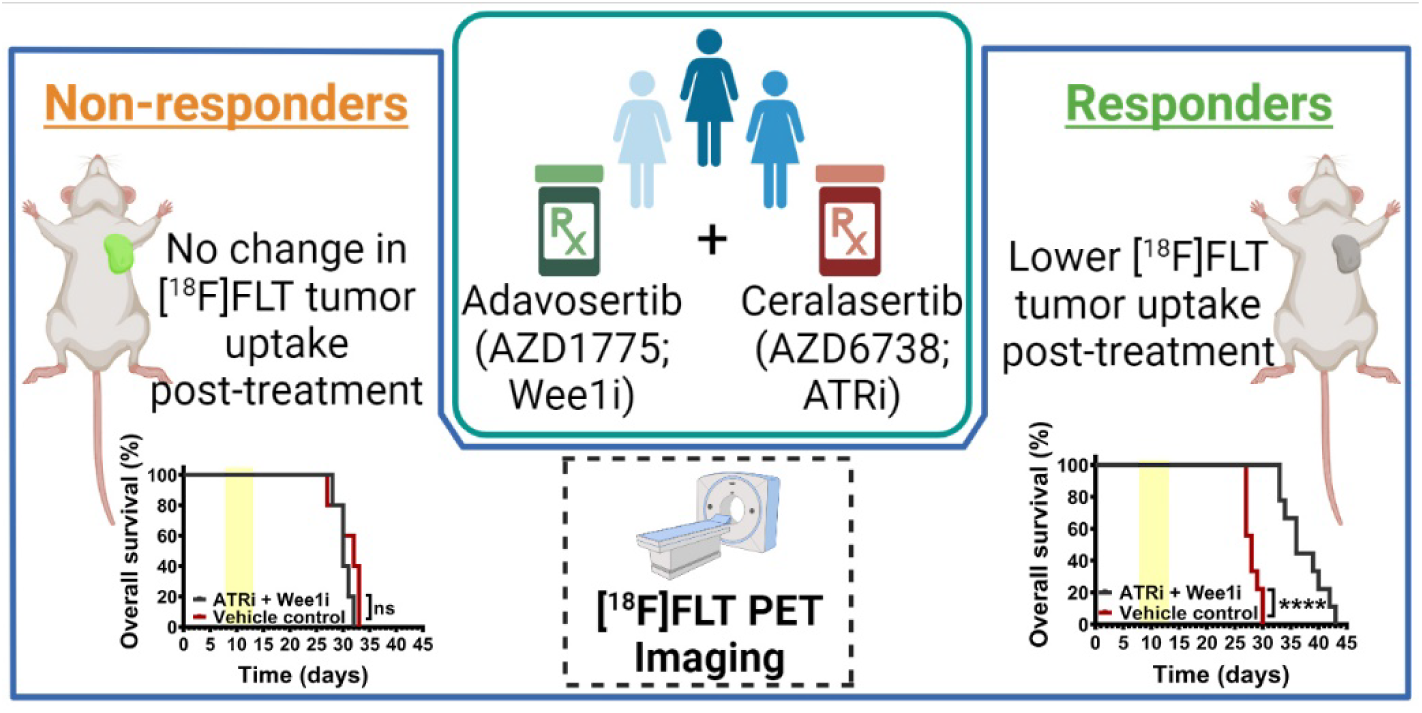

## Introduction

The clinical success of poly-ADP-ribose polymerase inhibitors, especially for cancers with deficiencies in homologous recombination, has heightened the interest in inhibitors of enzymes in the DNA damage response (DDR), particularly protein kinases involved in DNA damage signaling, as well as their combination with DNA damaging agents or cell cycle regulators. Recent preclinical and clinical studies have shown the efficacy of DDR inhibitors also in tumors with no known underlying genetic signatures, showcasing the urgency of novel biomarkers for treatment plans based on the principle of targeting the DDR. Of particular interest are non-invasive biomarkers that predict treatment response at an early stage of therapy.

Pre-clinical and clinical research aimed at evaluating chemotherapy response in neoadjuvant settings take advantage of combining imaging techniques (1) with histological findings to measure pathological response, identify potential prognostic biomarkers, and tailor personalized therapies. However, accurate measurement of treatment effectiveness to stratify cancer patients into responders and non-responders remains a challenge in the clinic. Early classification of non-responders may allow exploration of alternative therapies to avoid unnecessary side-effects of ineffective treatments. This requires reliable biomarkers that functionally detect and correlate early treatment-related changes in *tumor biology* with treatment outcome.

Non-invasive functional imaging of solid tumors utilizing positron emission tomography (PET) has become a widely used tool in the clinic (2, 3). Based on the radiotracer used, PET imaging provides semi-quantifiable data for radioactivity uptake in the form of standardized uptake values (SUV) as a measure of the specific underlying biological processes (4). PET-based imaging biomarkers allow for non-invasive longitudinal assessment of comprehensive spatial information across the entire tumor and body. PET with the glucose analog 2’-[^18^F]fluoro-2’-deoxyglucose (FDG) is widely used for metabolic imaging to sensitively detect malignant tumors (5, 6), but treatment induced inflammation can reduce the specificity of [^18^F]-FDG-PET (7–9).

To overcome this drawback and to measure cell proliferation status *in vivo*, [^18^F]3’-deoxy-3’-fluorothymidine (FLT) was used for PET (10). [^18^F]FLT cellular uptake is mediated *via* equilibrative nucleoside transporters 1 and 2 (ENT1/2) (11) and the phosphorylation of [^18^F]FLT by thymidine kinase 1 (TK1) traps the thymidine analogue inside the cell (12). Because TK1 is expressed during the S-phase of the cell cycle and remains inactive in quiescent cells (13, 14), [^18^F]FLT uptake was reported to be a surrogate marker for cell proliferation status (15, 16)(6). Several groups have reported a significant correlation between Ki-67 expression – the clinical gold standard for evaluating cell proliferation status – and [^18^F]FLT uptake in many different types of cancers (17–20), including those of the breast (4, 21, 22).

In the present pre-clinical study, we have evaluated changes in [^18^F]FLT tumor uptake as a non-invasive prognostic biomarker to identify early responders to combined AZD6738 (ATRi) and AZD1775 (Wee1i) treatment in a neoadjuvant therapeutic setting. AZD6738 and AZD1775 are bioavailable inhibitors of the kinases ataxia telangiectasia and Rad3 related (ATR) and Wee1. ATR is an apical kinase in the DDR, and Wee1 kinase regulates cell cycle progression. By combining inhibition of ATR and Wee1, we have recently shown cancer cell-specific synergistic cell killing which resulted in tumor control, metastasis inhibition, and increased overall survival of breast cancer xenograft bearing mice (23). Here, using a syngeneic orthotopic mouse model of triple-negative breast cancer (TNBC), we show that early post-treatment changes in [^18^F]FLT tumor uptake are promising predictive biomarkers for neoadjuvant chemotherapy with combined ATRi and Wee1i. Furthermore, using immunohistochemistry, we found a significant reduction in the percentage of Ki-67 positive cells following neoadjuvant chemotherapy with AZD6738 and AZD1775 in responding tumors, that correlated with the decrease in [^18^F]FLT uptake. Since AZD6738 and AZD1775 are currently being evaluated in clinical trials as monotherapy agents or in combination with various other genotoxic therapies, the present preclinical findings will have direct implications for future clinical trials to identify patient responders and non-responders to this combination treatment regime. While this study focuses on the combined inhibition of ATR and Wee1 as treatment strategy, we envision that changes in [^18^F]FLT uptake could be utilized also for other therapeutic strategies targeting the DDR and may become an integral part of an effort to use functional biomarkers to monitor therapy efficacy at an early stage for precision medicine.

## Methods

### Antibodies and chemicals

anti-Ki-67 (D3B5; #12202) was purchased from Cell Signalling Technologies. The bioavailable inhibitors AZD6738 and AZD1775 were kindly provided by AstraZeneca (Cambridge, UK).

### Cell lines

4T1 and EMT6 cell lines were purchased from the American Type Cell Culture (ATCC) and cultured in DMEM high glucose medium supplemented with 10% fetal calf serum. These cell lines were regularly tested for mycoplasma.

### Crystal violet assays

3000 cells per well were seeded into 96 well plates 4 h prior to drug treatment. Cells were treated with indicated concentrations of AZD6738 (100 nM to 4000 nM) and AZD1775 (50 nM to 2000 nM) for 96 h. After 96 h, cells were washed twice with 1X PBS and stained with 0.5% crystal violet (in 20% methanol) for 20 mins. Cells were then washed with water for 4 times, plates were air dried overnight. 200 μL methanol was added per well and incubated for 20 mins at room temperature. Optical density was measured at 584 nm using the FLUOstar Omega plate reader (BMG Labtech). Background values were subtracted using blank OD584. Data was calculated in terms of % surviving attached cells (% crystal violet OD) compared to vehicle control treated cells. Experiments were performed in triplicates in at least 3 independent experiments. Bliss cooperativity indices were calculated and plotted using the Combenefit software.

### Orthotopic breast cancer syngeneic mouse models and drug treatment

All animal experiments were carried out in accordance with guidelines of the Canadian Council on Animal Care and approved by the local animal care committee of the Cross Cancer Institute as protocol AC20251. Six weeks old female BALB/c mice were obtained from Charles Rivers Laboratories (Saint Constant, Québec, Canada). For tumor formation, 1 x 10^5^ 4T1 or 2 x 10^5^ EMT6 cells were mixed with Matrigel from Corning (Tewksbury, US) and PBS (1:1) and injected in 40 μL volumes orthotopically into the thoracic mammary fat pad of 8-10 week old female BALB/c mice. Tumor growth was measured every 3 days using a Vernier caliper and volumes were assessed as (length x width^2^)/2.

When tumors reached an approximate volume of 25-35 mm^3^ for 4T1 model or 60-80 mm^3^ for the EMT6 model, mice were randomly segregated into 2 groups (n = 9 per group) for each model. For consistency in tumor evolution, the time between inoculation and treatment was kept for both tumor models to 1 week. As EMT6 and 4T1 inoculation protocols were optimized for tumor take, this led to slightly different tumor volumes after 1 week. Mice were treated daily with vehicle or 25 mg/kg AZD6738 (in 10% DMSO, 40% polypropylene glycol, and 50% ddH_2_O) and 60 mg/kg AZD1775 (in 0.5% methylcellulose) combination *via* oral gavage for 5 days. Body weight was measured pre- and post-treatment as an indicator of toxicity. Mice were euthanized 12 hours post [^18^F]FLT-PET scans to allow for radioactivity decay and tumors were harvested for histology.

Treatment efficacy was categorized according to guidelines adapted from Response Evaluation Criteria in Solid Tumors (RECIST) (57, 58). The treatment efficacy was defined by the percentage change in tumor volume measured at the end of the treatment over the tumor volume measured before treatment by a Vernier calliper. Treatment response was classified as partial response (PR) if the reduction in total tumor size was > 30%; stable disease (SD) if the reduction in total tumor size was < 30% and tumor growth was < 20%; progressive disease (PD) if the growth in total tumor size exceeded 20% or new lesions were identified; and complete response (CR) if disappearance of tumor lesions was observed.

### Immunohistochemistry

Immunohistochemistry was performed on formalin fixed paraffin embedded (FFPE) tissue samples using standard procedures as previously described (3). Briefly, tumors were sectioned into 5 μm slices on precleaned Colorfrost Plus microscope slides (Fisher Scientific, USA) using a microtome (Leica, Germany). Tissue samples were baked at 60°C for 2 h and deparaffinized 3 times in xylene for 10 min each and subsequently rehydrated in a gradient of ethanol washes. For antigen retrieval, tissue sections were subjected to heat in a pressure cooker and 0.05% citraconic anhydride antigen retrieval buffer (pH – 7.4). Tissue samples were then blocked with 4% BSA for 30 min and incubated with the antibody against Ki-67 (D3B5; #12202; Cell Signaling Technologies; 1:500 dilution) overnight at 4°C. The next day, endogenous peroxidase activity was blocked for 30 min using 3% H_2_O_2_, followed by incubation with anti-rabbit HRP labelled secondary antibody (Dako EnVision+ System; K4007) for 1 h at room temperature in the dark. Samples were incubated with DAB (3,3’-diaminobenzidine) + substrate chromogen (Dako, USA) for brown color development, counterstained with hematoxylin, and mounted with DPX mounting medium (Sigma, USA). Images were captured using the Zeiss Axioskop2 plus upright microscope (Zeiss, Germany) equipped with Axiocam 512 color camera. At least 1000 cells were scored per sample.

### Radiosynthesis

[^18^F]3’-deoxy-3’-fluorothymidine (FLT) was synthesized by the method described by Machulla *et al.* (59) at the Cross Cancer Institute’s cyclotron facility and in house using a manual synthesis set up with 5’-O-(4,4’-dimethoxytrityl)-2,3’-anhydrothymidine (ABX GmbH, Radeberg, Germany) as the radiolabeling precursor.

### PET imaging

4T1 and EMT6 tumor-bearing BALB/c mice were anesthetized with isoflurane and placed on a 37°C heated bed to regulate their body temperature. 4-8 MBq [^18^F]FLT in 100-150 μL saline solution (4-8% ethanol) was injected through the lateral tail vein using a needle catheter. The presence of radioactivity in the injection solution was determined using a dose calibrator (AtomlabTM 300, Biodex Medical Systems, New York, NY, USA), which was cross-calibrated with the scanner. Mice were allowed to regain consciousness for about 40 min before anesthetizing them again and positioning in prone position into the centre of the field of view of an INVEON® PET scanner (Siemens Preclinical Solutions, Knoxville, TN, USA). A 10 min static PET scan was acquired at 60 min post-radioactivity injection in the three dimensions list mode. Imaging data were reconstructed using the maximum a *posteriori* (MAP) algorithm. The imaging data files were then further processed using the ROVER v2.0.51 software (ABX GmbH, Radeberg, Germany).

Masks defining the three dimensional regions of interest (ROI) over the tumor were defined and analyzed at a threshold of 50% of radioactivity uptake. Mean standardized uptake values were calculated for each ROI as SUV_mean_ = (measured radioactivity in the ROI / mL tumor tissue) / (total injected radioactivity / body weight of mouse).

### Statistical analysis

All statistical analysis was performed using the GraphPad Prism 8 (GraphPad Software, La Jolla, CA, USA). *P* values were calculated using one-way or two-way ANOVA (analysis of variance) tests and *P* < 0.05 was considered significant.

## Results

### Combined ATR and Wee1 inhibitor treatment leads to synergistic cell killing in 4T1, but not in the EMT6 murine breast cancer cell line

The serine/threonine kinase Wee1 regulates the inhibitory phosphorylation of CDK1 at tyrosine 15 to delay mitotic entry until suitable conditions have been met (24). Inhibition of Wee1 results in increased entry and prolonged mitosis making cancer cells more vulnerable to therapy induced mitotic catastrophe (25). ATR, the central kinase of the DDR to replication stress, controls cell cycle checkpoints by several pathways including the phosphorylation of the downstream kinase Chk1 in response to DNA damage. We recently showed that combined inhibition of ATR and Wee1 kinases results in synergistic cell killing in a variety of cancer cell lines (23). Our additional screening identified a murine breast cancer cell line, EMT6, that showed low ATR inhibitor sensitivity and did not display significant synergistic cell killing by ATR and Wee1 inhibition. As shown in **Fig. 1 A, C**, in a crystal violet assay with different concentrations of AZD6738 and AZD1775 for 4 days, the two murine breast cancer cell lines EMT6 and 4T1 show different sensitivities to the ATR inhibitor AZD6738. Furthermore, unlike in the 4T1 cell line (and over two dozen cell lines assayed previously (23)), ATR and Wee1 inhibition does not induce synergistic cell killing in EMT6 cells, as also confirmed by calculating Bliss combination indices (CIs) and synergy scores represented as a 3D matrix in **Fig. 1 B, D**. A CI value below 1 indicates synergy, and a value above 1 indicates antagonism **(Suppl. Table 1)**. The mechanisms behind the observed ATR inhibitor resistance and the absence of synthetic lethality of ATR and Wee1 inhibition in EMT6 remain to be investigated. While several groups have reported that cells with p53 and/or ATM defects are more sensitive to ATR inhibition (26–30) likely due to an impaired G1 checkpoint and/or increased replication stress due to relaxed S-phase entry (31), we did not see any correlation between p53 status and sensitivity to ATR inhibition in the human cancer cell lines tested in our previous study (23). We therefore suspect that the p53 wild type state of the EMT6 murine breast cancer cell line could be just one of several factors contributing to lower sensitivity to ATR inhibition **(Suppl. Table 2)**.

**Figure 1.**
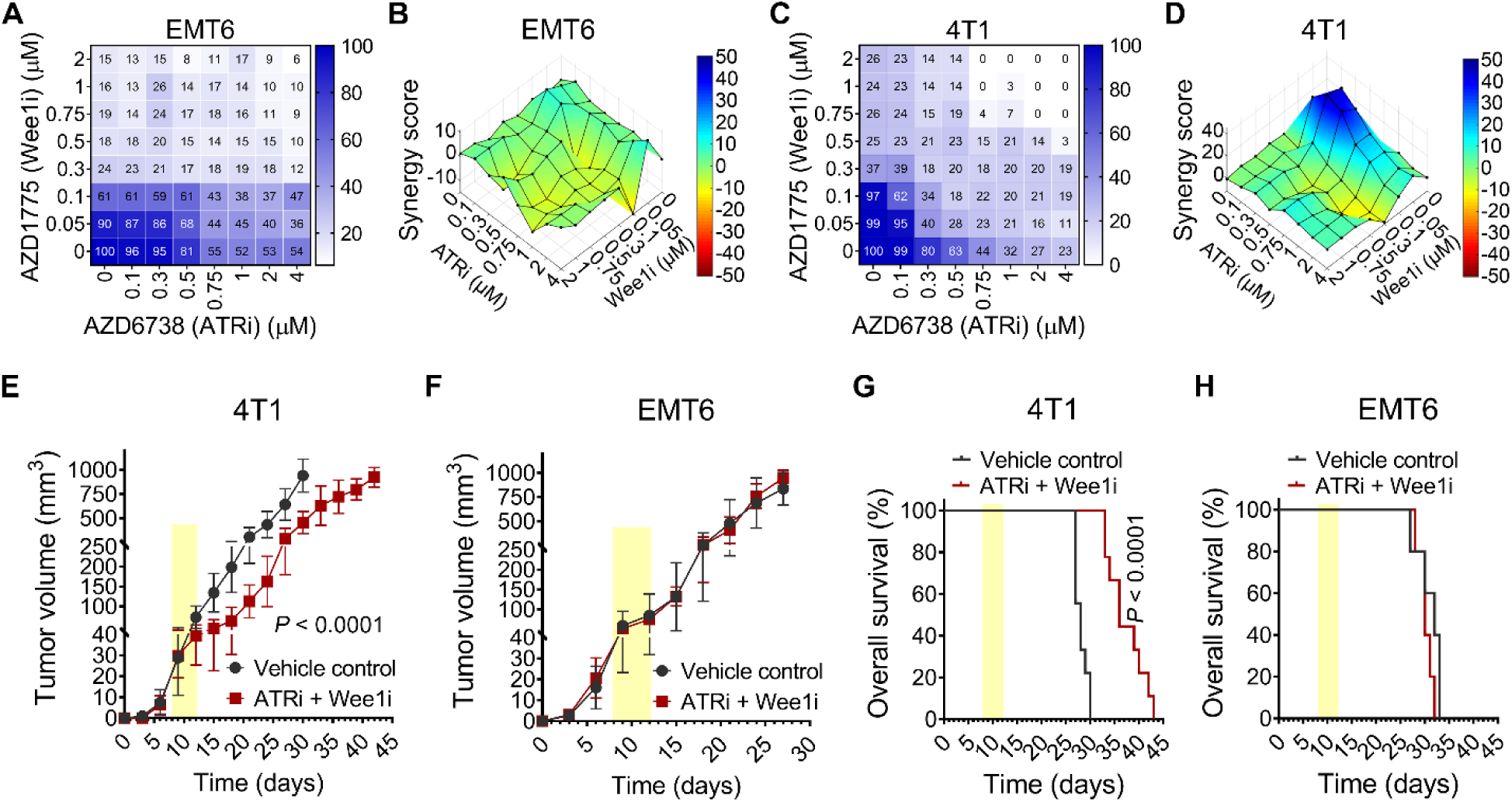
ATR and Wee1 inhibition leads to synergistic cell killing in 4T1, but not in EMT6, which is reflected by a difference in efficacy of the combination treatment *in vivo*. Murine breast cancer cell lines EMT6 **(A)** and 4T1 **(C)** were treated with increasing concentrations and different combinations of AZD6738 (up to 4 μM) and AZD1775 (up to 2 μM) for four days. Survival was assayed by crystal violet staining and experiments were repeated at least three times. The color bar indicates percent survival normalized to vehicle control-treated cells. **(B** and **D)** 3D plots were generated to represent the calculated synergy score in a 3D matrix. The blue color scale at low doses indicates strong synergy in 4T1 cells. No synergy was observed in EMT6 cells. **(E** and **F)** The graphs show tumor growth curves of 4T1 (n = 9 mice per group) and EMT6 (n = 5 mice per group) tumor bearing BALB/c mice treated with either vehicle or combined ATR and Wee1 inhibitors for 5 days (indicated by yellow shades). **(G** and **H)** Kaplan-Meier survival curves of the treated mice (n = 9 mice per group for 4T1 tumors, and n = 5 per mice group for EMT6 tumors).

The differing response of 4T1 and EMT6 to the drug combination treatment *in vitro* prompted us to use these specific cell lines as treatment responsive and refractory models in comparative *in vivo* studies. To investigate the drug efficacy and to test PET imaging as predictive non-invasive biomarker for treatment response we used an immune competent mouse model facilitated by the fact that 4T1 and EMT6 are both derived from tumors originating in BALB/c mice.

### EMT6 and 4T1 tumors as non-responder and responder models to combined ATR and Wee1 inhibition

To evaluate the therapeutic response of combined ATRi and Wee1i treatment *in vivo*, we established syngeneic orthotopic mouse breast tumors in BALB/c mice. The EMT6 and 4T1 allografts in BALB/c mice allowed for a robust evaluation of the drug response of metastatic TNBC by including an immune response, which was lacking in our previously analyzed xenograft models (23). Tumors were developed by injecting 4T1 or EMT6 cells **(Suppl. Table 2)** (32) in the thoracic mammary fat pad of 8-10 week old female BALB/c mice. In our previous study we had treated tumors for 25 days with the inhibitor combination (23). To validate [^18^F]FLT as a potential biomarker for *early* therapy response, during the present study mice were administered 25 mg/kg ATRi and 60 mg/kg Wee1i by oral gavage over just 5 days (days 1-5) and tumor growth kinetics monitored. As expected, 4T1 tumors in mice treated with combined ATRi and Wee1i for just 5 days showed a significantly growth delay in comparison to the vehicle control group (*P* < 0.001, two-way ANOVA) **(Fig. 1E)**. In contrast, no growth difference was observed between EMT6 tumors treated with drug combination or vehicle alone **(Fig. 1F)**. Furthermore, 4T1 tumor bearing mice treated with combined ATRi and Wee1i lived significantly longer despite the short treatment period of only 5 days (*P* < 0.01, log-rank Mantel-Cox test) – median survival of 41 days versus 28.5 days of the vehicle treated mice **(Fig. 1G)**. The overall survival of the EMT6 tumor bearing mice was similar in both the vehicle and combined ATRi and Wee1i treatment groups **(Fig. 1H)**. These observations not only were in line with the observed drug efficacies *in vitro*, but also validated the use of 4T1 and EMT6 tumors as models for responders and non-responders to combined ATR and Wee1 inhibition, even when administered for a short period.

### [^18^F]FLT-PET imaging as a predictive biomarker for combined ATR and Wee1 inhibitor therapy response in breast tumors *in vivo*

To validate [^18^F]FLT as a potential biomarker for *early* therapy response, mice were administered 25 mg/kg ATRi and 60 mg/kg Wee1i by oral gavage over just 5 days (days 1-5) and pre- (day 0) and post-treatment (day 7) PET scans were acquired to assess changes in [^18^F]FLT tumor uptake (**Fig. 2A)**. At least 5 mice were randomly selected from each group for further PET imaging a week later (day 14). Based on our *in vitro* findings, we expected EMT6 tumors to be poor responders to ATRi and Wee1i treatment *in vivo*. To validate this and to probe how [^18^F]FLT uptake correlated with tumor growth and treatment response, BALB/c mice bearing orthotopic EMT6 tumors were treated with ATRi and Wee1i as described above. Again, EMT6 tumors did not respond to the 5 days ATRi and Wee1i combination treatment regimen as no significant change in primary tumor volumes between the vehicle and treatment groups was measured on day 7 **(Fig. 2B)**. Of note, also the [^18^F]FLT tumor uptake measured either shortly after treatment (day 7; SUV_mean_ of 0.95 ± 0.10 for ATRi + Wee1i versus SUV_mean_ of 0.92 ± 0.06 for vehicle) or during the follow-up (day 14; SUV_mean_ of 0.96 ± 0.08 for ATRi + Wee1i versus SUV_mean_ of 0.90 ± 0.10 for vehicle) did not change significantly **(Fig. 2C, D)**, indicating that the mean [^18^F]FLT uptake did not change with tumor size over the observed period or as a result of the combination drug treatment. Representative maximum intensity projection PET images are shown in **Fig. 2C**.

**Figure 2.**
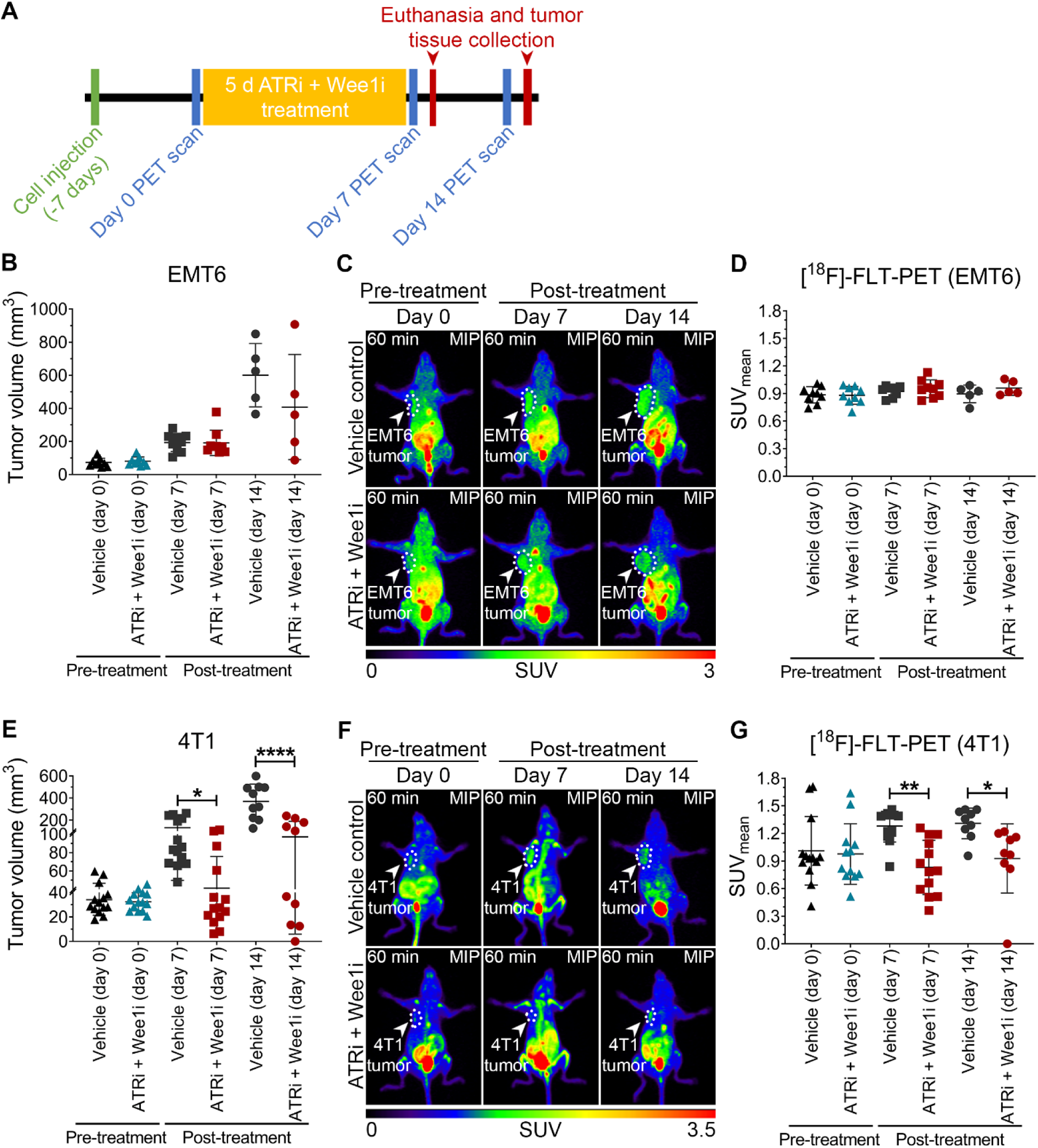
Combination treatment with ATR and Wee1 inhibitors and non-invasive monitoring of treatment response with [^18^F]FLT-PET imaging. BALB/c mice were injected orthotopically with 4T1 or EMT6 cells and treated for 5 days. Pre-and post-treatment [^18^F]FLT-PET scans were acquired to evaluate therapy response. **(A)** Schematic representation of the experimental outline. **(B** and **E)** Graphs represent pre- and post-treatment tumor growth in mice treated with ATRi + Wee1i combination or vehicle for 5 days. **(C** and **F)** Representative mouse images showing maximum intensity projection (MIP) of [^18^F]FLT uptake 60 minutes post radiotracer injection. **(D** and **G)** Quantitative [^18^F]FLT uptake shown as mean standardized uptake values (SUV_mean_). Data represented as scatter dot plot showing all experimental data points including mean values; error bars represent standard deviations. * indicates *P* < 0.05, ** indicates *P* < 0.01, **** indicates *P* < 0.0001 as calculated by one-way ANOVA.

We next performed the same experiments with 4T1 tumor bearing mice, which at the onset of treatment had a similar mean [^18^F]FLT tumor uptake (SUV_mean_; **Fig. 2G**). Interestingly, in the case of 4T1, 5 days of treatment with ATRi and Wee1i showed a significant reduction in tumor size on day 7 (*P* < 0.05; one-way ANOVA) and day 14 (*P* < 0.001; one-way ANOVA) **(Fig. 2E)**. Indeed, of the 9 mice followed longer, one animal even showed complete regression of the primary tumor (or complete response as per RECIST criteria) following just 5 days of the drug treatment as determined by the absence of palpable tumor at the primary site. Importantly, 4T1 tumors treated with ATRi and Wee1i had a significant reduction in [^18^F]FLT uptake compared to 4T1 tumors in control mice post-treatment (day 7), with SUV_mean_ of 1.28 ± 0.17 for tumors in vehicle control mice and SUV_mean_ of 0.83 ± 0.30 in treatment group mice (*P* < 0.01; one-way ANOVA; n = 14 mice per group) **(Fig. 2F, G)**. In 4T1 tumor bearing mice imaged at day 14, [^18^F]FLT tumor uptake continued to be significantly lower in mice treated with ATRi and Wee1i for just 5 days (SUV_mean_ of 0.93 ± 0.38) compared to mice receiving just the vehicle (SUV_mean_ of 1.31 ± 0.17) (*P* < 0.05; one-way ANOVA; n = 9 mice per group) **(Fig. 2G)**.

### Early changes in [^18^F]FLT uptake correlate with treatment response

To evaluate treatment response, we adapted the Response Evaluation Criteria in Solid Tumors (RECIST) that is routinely used in the clinic to define treatment efficacy. While the clinical assessment of RECIST uses tumor volume estimates based on CT scans, here we relied on measurements using a caliper, which is a standard method in pre-clinical studies. Based on this, we found that 5 out of 14 (∼35%) 4T1 tumor-bearing mice showed PR (indicated in blue) to combined ATRi and Wee1i treatment for just 5 days and 5 (∼35%) presented with SD (indicated in orange) on day 7. Interestingly, 2 out of 9 (∼22%) 4T1 tumor bearing mice continued to show PR even one week after treatment completion (day 14) and 1 out of 9 (∼11%) had CR (indicated in green) **(Fig. 3A)**. These responses were in line with the early changes detected with [^18^F]FLT PET scans performed after treatment completion. In contrast, 9 out of 9 (100%) EMT6 tumor bearing mice indicated PD (indicated in black) at day 7 measurements and 4 out of 5 (80%) continued to show PD with one mouse (20%) showing indication of SD on day 14 **(Fig. 3B)**.

**Figure 3.**
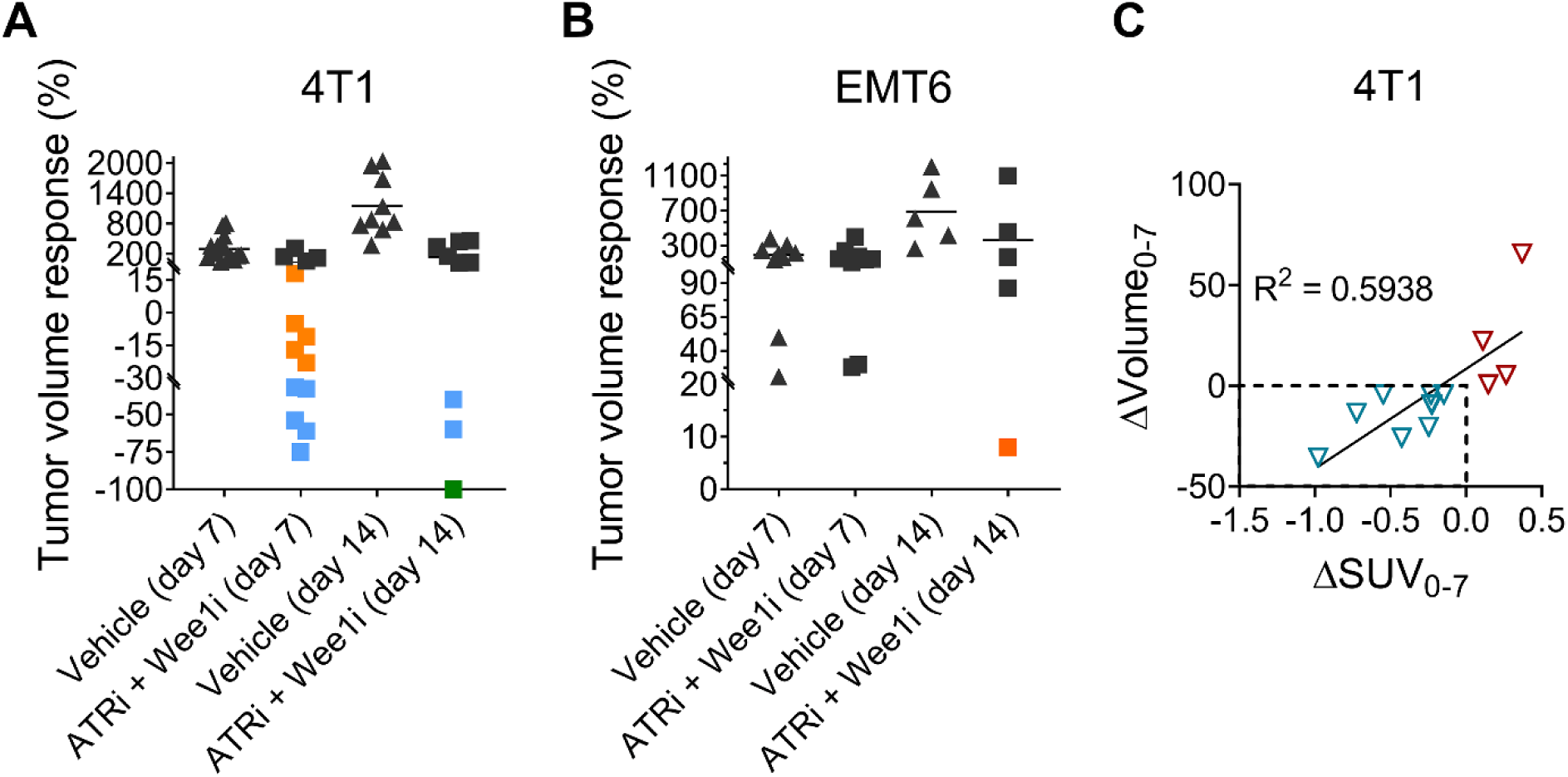
Effect of ATR and Wee1 inhibitors on tumor volume response. Graphs represent tumor volume response calculated based on adapted RECIST criteria for 4T1 **(A)** and EMT6 **(B)** tumor bearing mice treated with ATRi and Wee1i for 5 days. Disappearance of all lesions indicates complete response (CR; indicated in green); tumor volume reduction > 30% indicates partial response (PR; indicated in blue) and < 30% stable disease (SD; indicated in orange), whereas growth > 20% is classified as progressive disease (PD; indicated in **black**) (Notice the difference in the y-axis). **(C)** Difference in SUV_mean_ correlates with changes in tumor volume for individual 4T1 tumors. The graph shows change in SUV_mean_ (ΔSUV_mean_) versus change in tumor volume (ΔVolume) post-treatment with combined ATR and Wee1 inhibitors for 5 days. A decrease in [^18^F]FLT uptake after 7 days correlated with reduced tumor volume (indicated in teal) whereas no tumor shrinkage was seen in tumors where the ΔSUV_mean_ did not decrease (indicated in red).

### [^18^F]FLT uptake correlates with Ki-67 immunohistochemical staining

Although an indirect biomarker for proliferation, Ki-67 has become the “gold standard” for measuring cell proliferation status of biopsy samples in the clinic. Recently, Ki-67 was reported to be constitutively expressed during the S, G2, and M phases of the cell cycle, whereas – while Ki-67 expression during the G0/quiescent and G1 phases is generally lower – the dynamics of Ki-67 levels in non-proliferating cells vary (33). [^18^F]FLT uptake was originally proposed as an imaging biomarker for cell proliferation status based on its entrapment by TK1 in the thymidine salvage pathway (10). Nevertheless, the correlation between [^18^F]FLT uptake and TK1 levels or activity or Ki-67 levels remains unclear. Several contradictory findings reported indicate a more complex relationship between these parameters (34). To correlate [^18^F]FLT uptake with Ki-67 staining in our *in vivo* experiments, we have performed immunohistochemistry with slices from 4T1 **(Fig. 4A)** and EMT6 **(Fig. 4D)** tumors collected post-imaging. Tumor tissues were excised on day 8, 12 hours after [^18^F]FLT-PET scans from vehicle control mice and mice treated with the ATRi and Wee1i combination for 5 days (n = 4 mice per group). In support of a correlation of [^18^F]FLT PET uptake with proliferation *in vivo* (R^2^ = 0.7459), we observed a significant reduction in Ki-67 positive cells in 4T1 tumors treated with ATRi and Wee1i for just 5 days (*P* < 0.05; two-way ANOVA) compared to control tumors **(Fig. 4A-C)**. In contrast, no differences were found in Ki-67 staining between vehicle or ATRi and Wee1i treated EMT6 tumors **(Fig. 4D, E)**.

**Figure 4.**
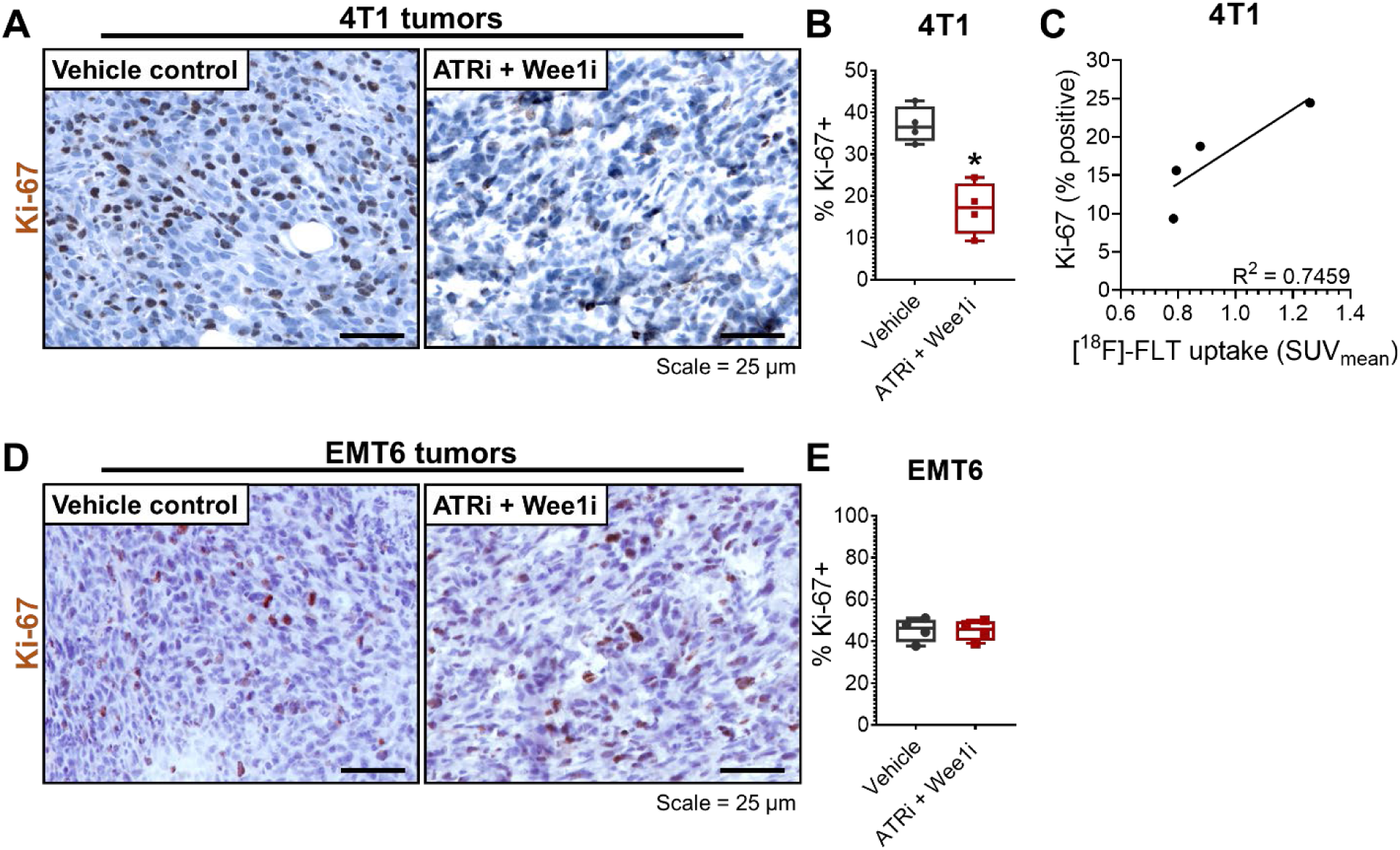
Histological evaluation of cell proliferation status. Tumors treated for 5 day with ATRi + Wee1i or vehicle control were harvested 12 hours after [^18^F]FLT-PET scan and processed for immunohistochemistry. **(A** and **D)** Representative images of the 4T1 and EMT6 tumor sections stained with Ki-67 marker to evaluate cell proliferation status. **(B** and **E)** Graphs show percentage of Ki-67 positive cells. **(C)** The graph plots the correlation between Ki-67 staining and [^18^F]FLT uptake. Data represent evaluation from 4 independent tumors in each group. * indicates *P* < 0.05 (two-way ANOVA).

## Discussion

AZD6738 and AZD1775 are currently being evaluated in Phase I/II clinical trials in combination with several other genotoxic agents (35). In addition, preclinical studies of AZD6738 or AZD1775 as radiosensitizers might lead to their use as adjuvant to radiation therapy (36–42). Therefore, identifying ways to accurately predict treatment response and classify patients as responders or non-responders to AZD6738/AZD1775 (or other therapies targeting the DDR) at an early stage of clinical treatment has the potential to greatly impact cancer therapy, by avoiding unnecessary therapy side effects and helping clinicians opt for alternative treatment options. Molecular imaging of tumors provides excellent alternatives to visualize treatment outcomes non-invasively. In that regard, functional PET imaging is a particularly attractive approach due to its ability to assess molecular changes within tumors before size related physical changes manifest.

### EMT6 as model cell line for cancers refractory to ATR and Wee1 inhibitor treatment

TNBC is often invasive and very aggressive (43). Using a panel of TNBC cells, we identified EMT6 as resistant to ATR inhibition compared to other cell lines, including 4T1. EMT6 was also the only cell line we found so far in which the ATR and Wee1 inhibitor combination failed to show synergistic cell killing *in vitro*. In stark contrast to 4T1 tumors, the combination therapy was also ineffective for EMT6 tumors *in vivo*. The underlying mechanisms for EMT6 resistance remain unclear, yet a recent a genome-wide CRISPR knockout screen showed that individual inactivation of seven genes (KNTC1, EEF1B2, LUC7L3, SOD2, MED12, RETSAT, and LIAS) promoted resistance to the ATR inhibitors AZD6738 and M6620 (44). While the status of these genes in the murine TNBC lines EMT6 and 4T1 is currently unknown, these genes may serve as a starting point for future studies evaluating drug resistance mechanisms to combined ATR and Wee1 inhibition. Yet the reported screen entirely relied on single gene knockout. It is likely that multifactorial genetic changes during carcinogenesis and associated changes in expression levels, not necessarily complete loss, of genes are determining drug resistance. As an example, we previously identified increased expression of the kinase MYT1 as a mechanism of acquired Wee1 inhibitor resistance (45).

#### Changes in [^18^F]FLT uptake during early treatment stage as biomarker for tumor response

Our pre-clinical results indicate that early changes in [^18^F]FLT tumor uptake following just 5 days of combined ATR and Wee1 inhibitor treatment may be used to predict responders to this therapy. Using an adapted RECIST criteria, we were able to correlate the initial reduction in radiotracer uptake to therapy response as the trend in [^18^F]FLT uptake a) correctly identified 4T1 as treatment-sensitive tumors compared to EMT6 and b) was able to predict the 4 out of 14 individual 4T1 tumors that did not respond favourably to the drug combination. Importantly, [^18^F]FLT uptake *at the onset* of treatment was unable to predict treatment outcome, as both tumor types, EMT6 and 4T1, showed similar [^18^F]FLT uptake pre-treatment. SUVs for [^18^F]FLT in both, 4T1 and EMT6 tumors, measured around 0.9 at the onset of treatment, but EMT6 failed to respond to the drug combination (**Fig. 1F,H**). Consequently, only the *change* in [^18^F]FLT tumor uptake over the short treatment period could serve as predictive biomarker for the ATR and Wee1 inhibition efficacy. A further analysis reveals a quantitative correlation between the shift in SUV (ΔSUV_0-7_) and the change in tumor volume (ΔVolume_0-7_) at day 7 (R^2^ = 0.5938) **(Fig. 3C)** for the individual 4T1 tumors. Indeed, only the tumors that showed a decreased [^18^F]FLT uptake post-treatment (at day 7), also showed a reduction in tumor volume due to the 5 day combined ATRi and Wee1i treatment. Noteworthy, three 4T1 tumors that showed just a reduction in the growth rate compared to untreated tumors, but no tumor shrinkage, did not display a decrease in [^18^F]FLT uptake.

Changes in [^18^F]FLT-uptake have been used previously, including at early stages of treatment, to predict tumor response for cytotoxic chemotherapy, such as anthracycline, taxane or platinum-based regimens (46–48), or radiation therapy (49, 50). [^18^F]FLT-PET has also been proposed for monitoring cytostatic treatments, such as with mTOR (51) or epidermal growth factor (EGFR) inhibitors (52). Yet to our knowledge the present study reports for the first time that [^18^F]FLT-PET could be used as predictive biomarker for treatment with inhibitors of the DDR.

#### Inhibiting the DDR, [^18^F]FLT-uptake and Ki-67 staining

As we showed previously, combined ATR and Wee1 inhibition leads to cancer cell death due to untimely mitotic entry (23). The unrepaired DNA damage or under-replicated DNA leading to mitotic catastrophe after G2/M checkpoint abrogation is in large part due to the high replication stress found in cancer cells due to their frequent loss of the G1/S checkpoint, polyploidy and/or genome instability. It is therefore expected that a higher proliferative index would correlate with a higher percentage of cells with replication stress and thus with susceptibility to DDR inhibitors. [^18^F]FLT-uptake is often considered an indicator of the proliferative state of tumors and we were therefore surprised that [^18^F]FLT-uptake at the onset of ATR/Wee1 inhibitor treatment was unable to predict tumor response. Previous breast cancer studies reported a positive correlation between [^18^F]FLT-PET uptake and Ki-67 staining, the standard immunohistochemistry biomarker for proliferative tissues in the clinic (4, 21, 53, 54). Yet due to contradictory reports (55), we corroborated the correlation in our mouse model **(Fig. 4)**. The implications are that the proliferative state of the tumor pre-treatment is unsuited to predict the response to inhibitors of ATR and Wee1 (and by extension inhibitors of the G2/M checkpoint or DNA damage inhibitors in general). Possible explanations are differences in DNA repair capacity or thresholds in checkpoint activation, alternative pathways to prevent entry into mitosis (such as upregulation of the kinase MYT1 (45)), or differential cellular drug uptake.

Yet the observed predictive power of changes in [^18^F]FLT-uptake over a short period of combination drug therapy is very promising for identifying patients responding to ATR/Wee1 inhibition and thus allowing the personalization of treatment plans. Future studies could show whether [^18^F]FLT-PET due to its non-invasiveness (and high specificity compared to [^18^F]FDG-PET (56)), can become a valuable clinical biomarker also for other inhibitors of the DDR. That class of inhibitors is currently one of the most promising contributors to improved cancer therapy and a range of drugs with new targets in the DDR are entering clinical trials and practice.

### Author contributions

ABB, MW, FW, and AMG designed the experiments. ABB performed most experiments, analyzed data, and performed statistical analysis. MW performed PET scans and analyzed PET imaging data. FW and AMG conceived the data. ABB and AMG wrote the manuscript.

## Acknowledgments

The authors thank Floyd Baker, David Clendening, and Blake Lazurko from the Edmonton Radiopharmaceutical Center for ^18^F production on a TR-19 cyclotron (Advanced Cyclotron Systems Inc, Vancouver, BC, Canada); Cody Bergman (Department of Oncology, University of Alberta) for performing the radiosynthesis of [^18^F]FLT; Dan McGinn (Cross Cancer Institute) for support with animal work; Dr. Hans-Soenke Jans (Department of Oncology, University of Alberta) for technical support of the preclinical PET imaging facility; and AstraZeneca for compounds AZD6738 and AZD1775. ABB is supported by Alberta Cancer Foundation’s Dr. Cyril M. Kay Graduate Scholarship. This research is funded by a grant from the Canadian Institute of Health Research (AMG) and the Dianne and Irving Kipnes Foundation (FW).

