## Supplementary Information for "[^18^F]FLT-PET as predictive non-invasive biomarker for neoadjuvant therapy with Wee1 and ATR inhibitors"

| *Cell Line* | *AZD6738 (ATRi)* | *AZD1775 (Wee1i)* | *Bliss CI* | *ATRi $IC_{50}$* | *Wee1i $IC_{50}$* |
|---|---|---|---|---|---|
| ***4T1*** | 100 nM | 50 nM | 0.39 | 736 nM | 308 nM |
| | 100 nM | 100 nM | 0.10 | | |
| | 300 nM | 50 nM | 0.34 | | |
| | 300 nM | 100 nM | 0.33 | | |
| ***EMT6*** | 100 nM | 50 nM | 1.04 | 2578 nM | 162 nM |
| | 100 nM | 100 nM | 1.06 | | |
| | 300 nM | 50 nM | 1.03 | | |
| | 300 nM | 100 nM | 1.02 | | |

**Suppl. Table 1. Synergistic murine breast cancer cell killing by ATR and Wee1 inhibition.**
$IC_{50}$ values and Bliss combination indices (CI) at indicated drug concentrations calculated from at least three independent experiments. A Bliss CI of less than 1 indicates drug synergy, a CI of < 0.7 strong synergy, and a CI of 1 pure additivity.

| *Gene* | *4T1* | *EMT6* |
|---|---|---|
| ***TP53*** | Mutant | Wildtype |
| ***ATM*** | Wildtype | Wildtype |
| ***BRCA1*** | Wildtype | Wildtype |
| ***BRCA2*** | Wildtype | Wildtype |
| ***PIK3CG*** | Mutant | ? |
| ***PTEN*** | Wildtype | Mutant (G209*) |

**Suppl. Table 2. Known mutation status in 4T1 and EMT6 murine breast cancer cell lines of genes with potential impact on ATR inhibitor sensitivity.**
Data courtesy of Charles River.
